# Oscillating Ears Dynamically Transform Echoes in Constant-Frequency Bats

**DOI:** 10.1101/2025.06.14.659613

**Authors:** Ravi Umadi

## Abstract

Constant-frequency (CF) bats exhibit rapid oscillations of their external ears. Yet, the functional role of these movements has remained unresolved since their initial documentation over half a century ago. Although recent studies have demonstrated that pinna motion generates Doppler shifts, they do not explain why ear oscillations intensify at close range or how these dynamics contribute to echo perception. In this study, I investigate the hypothesis that oscillatory ear movements enhance echo information during CF echolocation. Using a simplified receiver-motion model, I examine how time-varying pinna pose reshapes the temporal and spectral structure of returning echoes. I show that ear oscillations inject dynamic transformations into the received signal, producing multiple informative views of the same echo and increasing both temporal contrast and spectral diversity around the CF carrier. These transformations are strongest under behavioural conditions in which target-state uncertainty is expected to be high, offering a potential functional explanation for the long-standing observation that ear-oscillation rate increases as bats approach a target. The results suggest that oscillatory ear movements act as an adaptive, receiver-side mechanism that enhances echo information during CF echolocation, complementing the well-known emitter-side adaptations of high-duty-cycle biosonar.

## 1 INTRODUCTION

Bats belonging to the families *Rhinolophidae* and *Hipposideridae* rely on long, narrowband biosonar pulses (Constant Frequency (CF) calls) whose echoes are analysed with exceptional frequency resolution. This acoustic specialisation enables the detection of minute Doppler shifts and underlies the well-known mechanism of Doppler-shift compensation, whereby bats adjust their emission frequency so that returning echoes remain centred on the auditory fovea [1]. However, this dependence on narrowband signals imposes a constraint: CF echoes contain relatively little intrinsic temporal or spectral structure. This limitation becomes most apparent during close-range hunting, where pulse durations shorten, pulse rates increase, and echo delays collapse to a few milliseconds. Under these conditions, the outgoing call and incoming echo frequently overlap in time and frequency. Because relative velocity decreases as the bat nears its target, classical Doppler shifts become too small to segregate call and echo, raising the fundamental question of how CF bats preserve reliable echo information during the most demanding phases of echolocation.

A distinctive feature of CF bats is their rapid and rhythmic oscillation of the external ears. Schneider and Möhres (1960) [2] first demonstrated that immobilising the pinna musculature severely disrupts orientation, forcing bats to compensate with vigorous head movements. Subsequent studies by Griffin et. al. (1962) [3] and Pye and Roberts (1970)[4] revealed that ear oscillations are tightly but not perfectly coupled to sonar pulse emission, and — critically — that oscillation frequency increases markedly as bats approach prey. Pye and Roberts reported oscillation rates of 30–35 Hz with angular excursions of ~ 30° and tip velocities of roughly 30 cm/s. At CF frequencies of 50–100 kHz, such local motion would generate instantaneous Doppler shifts of 4–9 kHz (0.12 octaves) if applied uniformly across the ear, and even the velocity gradients along the deformable pinna are sufficient to introduce detectable frequency perturbations.

Modern work by Yin and Müller (2019) [5] confirmed that pinna motion generates Doppler components above psychophysical thresholds and that these components enter the ear canal. Their biomimetic model demonstrated direction-dependent spectral signatures created by moving ear surfaces, establishing that oscillatory reception can impose measurable acoustic transformations. However, these findings do not explain two long-standing behavioural observations. First, ear oscillation rates increase specifically during close-range, high-demand conditions in which call–echo overlap is strongest. Second, CF bats cannot successfully hunt without oscillating their ears, despite possessing highly refined spectral sensitivity. Existing models do not address why these oscillations are behaviourally indispensable or how they contribute to perception when Doppler-based segregation of call and echo becomes unreliable.

Long CF pulses pose an additional challenge: echoes often return while the bat is still emitting. At echo delays below ~5–10 ms, the returning signal overlaps the forward-emitted call, and because the bat’s approach velocity declines near the target, Doppler-shift compensation cannot separate the two signals. The outgoing call remains spectrally stable, whereas the structure of the incoming echo becomes increasingly difficult to extract due to masking and frequency overlap. Classical Doppler theory [6] predicts that oscillatory receiver motion can impose timevarying frequency and phase modulations on an incident sound. Thus, the moving ears could impart transformations on the *echo only*, creating spectral and temporal features that are absent from the outgoing call. This suggests a potential mechanism by which ear oscillations help the bat segregate call from echo during overlap, even when frequency separation is small.

Building on this premise, and on the early proposal by Pye and Roberts [4] that ear movements play an active role during echo reception, I propose that oscillatory pinna motion constitutes an adaptive, receiver-side mechanism for enhancing echo information during constant-frequency echolocation. Specifically, I hypothesise that increasing ear-oscillation frequency at close range allows CF bats to compress multiple cycles of binaural contrast into an increasingly short echo-analysis window as call duration and echo delay decrease.

Because the ears oscillate in antiphase, each ear samples the incoming echo with a distinct, continuously changing velocity profile and orientation. This produces echo-specific instantaneous frequency modulations, amplitude fluctuations, and spectral sidebands — dynamic features that are absent from the outgoing call and therefore remain identifiable despite call–echo overlap. Spectral broadening introduced by oscillatory motion may thus enrich the inherently sparse CF echo by adding sidebands that increase spectral contrast, improve sensitivity to small target-induced frequency shifts (e.g. wingbeat micro-Doppler), and enhance binaural disparities that support directional discrimination. In this view, spectral broadening is not a secondary by-product of ear motion but a beneficial and biologically exploited transformation that boosts perceptual salience under conditions where conventional Doppler mechanisms fail.

To investigate this hypothesis, I developed a computational model in which each ear is represented as a 2 cm lever oscillating sinusoidally in antiphase at 5–50 Hz with a maximum deflection of 45°. Echoes were synthesised as the sum of locally Doppler-shifted components arising from segments along the ear, combined with a baseline Doppler shift due to forward flight. Instantaneous frequency and amplitude were estimated using the analytic signal. The simulations revealed that oscillatory reception produces fluctuations in instantaneous frequency exceeding 0.5–1 kHz over a single oscillatory cycle and generates substantial spectral broadening beyond that predicted by average velocity alone. These transformations were not mirror opposites across ears but instead reflected the phase-offset kinematics of natural pinna motion. To provide an empirical analogue, I recorded reflections of an 80 kHz tone from a sinusoidally oscillated subwoofer. The spectral spread increased from ~234 Hz in static conditions to over 1.2 kHz at 30 Hz oscillation, confirming that oscillatory motion produces non-linear, biologically relevant modulations of narrowband signals.

Together, these findings support a functional interpretation of ear oscillations in CF bats: by dynamically reshaping the spectral and temporal structure of echoes, oscillating ears enhance echo information precisely when perceptual demands are highest. This framework explains why ear oscillation rates increase with decreasing target range, why CF bats cannot hunt without these movements, and how oscillatory reception may help resolve call–echo overlap when classical Doppler cues are insufficient. In this view, ear motion is not merely an accessory behaviour but a critical component of the biosonar system that expands the informational content of returning echoes during constant-frequency echolocation.

## 2. METHODS

The aim of the present methods is to formalise and test a receiver-side mechanism by which oscillatory ear motion (Fig. 1) can transform the structure of CF echoes under conditions of call–echo overlap. To this end, I developed a theoretical framework that treats the bat’s pinnae as dynamically moving receivers whose time-varying kinematics impose Doppler- and phase-based transformations on incoming echoes without affecting the emitted signal. This framework explicitly distinguishes between Doppler shifts arising from linear flight and object motion, and modulation effects generated locally by oscillatory ear motion. It further incorporates the antiphase coupling of left and right ears observed in CF bats, allowing binaural disparities to emerge naturally from receiver dynamics.

**Figure 1:**
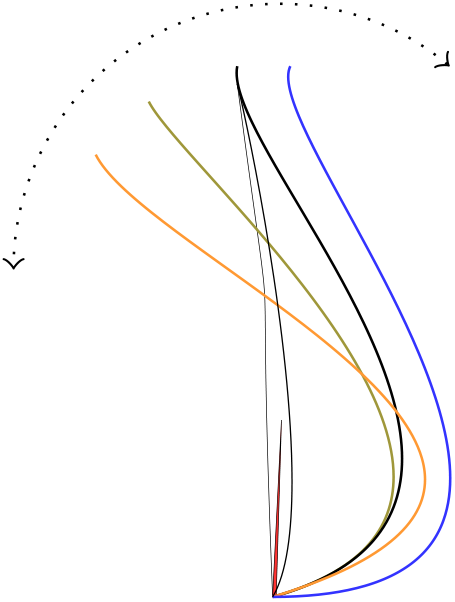
Mechanism of Doppler-induced signal transformation across an oscillating bat ear. As an echo impinges on the bat’s ear, it reaches different regions of the ear surface with a slight delay due to the ear’s spatial extent. Simultaneously, different points along the ear move at different velocities due to the ear’s oscillatory motion. This combination of delayed arrival and spatially varying velocity results in a gradient of instantaneous Doppler shifts. Each point along the ear effectively receives a frequency-shifted version of the echo, with the tip of the ear contributing the largest shifts due to its highest velocity. This dynamic velocity profile imparts a time-varying phase transformation across the reflected wavefront, causing phase warping and the generation of interference patterns in the time domain. In the frequency domain, these effects appear as a spread of energy around the carrier frequency, resulting in a widened spectral band. This phenomenon is distinct from the classical Doppler shift induced by linear relative motion, which produces only a uniform frequency offset.

Building on this analytical foundation, I implemented a computational model that represents each ear as a simplified oscillating structure with a distributed velocity field along its length. This abstraction is intentionally minimal: it does not attempt to reproduce detailed pinna morphology or static head-related transfer functions, but instead isolates the acoustic consequences of oscillatory receiver motion per se. The simulations quantify how ear kinematics shape instantaneous frequency, spectral bandwidth, phase structure, and binaural differences in received echoes across a biologically relevant parameter space.

Finally, to validate the underlying acoustic principle independently of anatomical detail, I conducted a controlled physical experiment in which an ultrasonic tone was reflected from a sinusoidally oscillating surface. This experimental analogue tests the same receiver-motion mechanism predicted by the theory and simulations, demonstrating that oscillatory motion alone is sufficient to induce nonlinear spectral broadening and phase warping of narrowband signals. Together, these methods establish a coherent progression from theoretical formulation to numerical prediction and empirical validation of oscillatory reception in CF echolocation.

### 2.1 Theoretical Framework for Oscillatory-Receiver Doppler Modulation

#### 2.1.1 Kinematics and Notation

I represent the bat as a moving emitter–receiver in a three-dimensional environment containing *N* reflecting objects. The bat’s body is at position **r**_b_(*t*) with velocity **v**_b_(*t*), and object *i* is at position **r**_*i*_(*t*) with velocity **v**_*i*_(*t*), *i* = 1, …, *N*. The unit vector from the bat to object *i* is

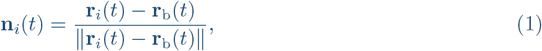

and the radial relative velocity between bat and object *i* is

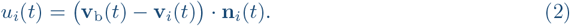

The emitted call is a narrowband CF signal with instantaneous frequency *f*_em_(*t*), centred around a resting frequency *f*_0_ in the absence of Doppler-shift compensation (DSC). Sound propagates at speed *c*.

For simplicity, I treat each reflector as a point scatterer with scalar reflectivity *A*_*i*_ and assume specular echoes; extensions to extended reflectors do not change the core argument.

The one-way propagation delay from bat to object *i* at emission time *t* is *τ*_*i*_(*t*) ≈ *d*_*i*_(*t*)*/c*, where *d*_*i*_(*t*) = ∥**r**_*i*_(*t*) − **r**_b_(*t*)∥ is the distance at emission. The echo returns at time *t* + *τ*_*i*_(*t*).

#### 2.1.2 Doppler Shifts from Bat and Objects

For small velocities relative to *c*, the dominant Doppler shift for a pulse emitted at frequency *f*_em_ and reflected by object *i* can be approximated from the radial relative velocity *u*_*i*_ as

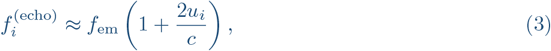

where the factor of 2 accounts for the bat acting as both moving source and moving receiver with respect to the reflector.

For a stationary object (**v**_*i*_ = **0**), the shift depends only on the bat’s own velocity component along **n**_*i*_. For a flying insect, **v**_*i*_ introduces an additional, independent contribution.

#### 2.1.3 Two Doppler-shift Compensation Strategies

I distinguish two idealised DSC strategies.

##### Target-specific DSC

If the bat were to lock onto a particular object *k* and use it as a reference, it would adjust the emission frequency *f*_em_(*t*) such that the echo from that object is centred on a preferred reference frequency *f*_ref_:

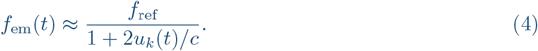

The echo from object *k* then satisfies

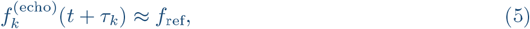

whereas echoes from all other objects *i* ≠ *k* are shifted according to their own relative velocities *u*_*i*_.

##### Global-reference DSC

Behavioural data for *Rhinolophus ferrumequinum* indicate that CF bats do not compensate for Doppler shifts created by moving insects, but stabilise echoes from the stationary environment instead [1]. This corresponds to a global-reference strategy in which *f*_em_ is adjusted to keep the average Doppler shift from quasi-static reflectors (ground, walls, foliage) at *f*_ref_.

Formally, let 𝒢 denote the set of “global” stationary reflectors (e.g. environment) with *u*_*i*_ ≈ **v**_b_ · **n**_*i*_. The bat adjusts *f*_em_(*t*) such that

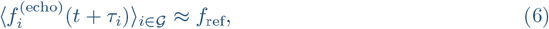

which, using Eq. (3), gives the approximate control law

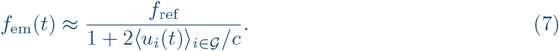

Echoes from moving insects then appear at

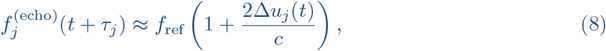

where Δ*u*_*j*_ = *u*_*j*_ −⟨*u*_*i*_⟩_*i∈ℊ*_ represents the insect’s relative motion with respect to the background.

### 2.2 Feasibility of Target-specific DSC During Foraging

In a natural foraging environment the bat encounters a large, time-varying set of reflectors:

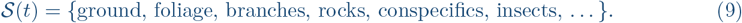

Each element *i* ∈ 𝒮 has its own radial velocity *u*_*i*_(*t*) and contributes a superposed echo. For target-specific DSC to operate, the bat would need to (i) identify a single reflector *k* in this cluttered stream, (ii) estimate its instantaneous *u*_*k*_(*t*), and (iii) adjust *f*_em_(*t*) on a call-by-call basis according to Eq. (4).

However, behavioural evidence shows that CF bats do not compensate for insect motion [1]. Instead, echoes from moving targets retain their Doppler structure relative to a globally stabilised background. This suggests that, during general foraging without a locked target, the bat’s DSC system operates according to Eq. (7), using quasi-stationary environmental echoes as a global reference.

Under this global strategy, the bat receives a dense stream of echoes, many of which originate from objects that are not yet behaviourally selected. Distinguishing and localising potential targets in this stream is therefore non-trivial, particularly because CF pulses are long and narrowband, and call–echo overlap is common at short ranges.

#### 2.2.1 Oscillating Ears as Dynamic Receivers

I now incorporate oscillatory ear motion into the model. The left and right ears are separated by an interaural baseline *b* and oscillate in antiphase with angular displacement

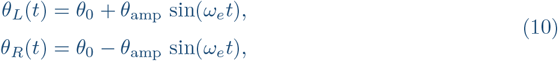

where *θ*_0_ is the mean pinna angle, *θ*_amp_ the oscillation amplitude, and *ω*_*e*_ = 2*πf*_*e*_ the ear oscillation angular frequency. For clarity, I treat each ear as a rigid lever of length *L* pivoting at its base; more realistic elastic deformations will only enhance the effects described below.

The linear velocity of the ear tip (or another representative point on the pinna surface) is

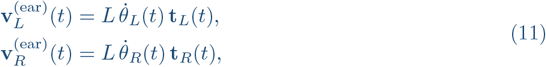

where **t**_*L,R*_(*t*) are unit tangent vectors along the arc of motion. For small angles and a fixed rotation axis, 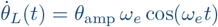 and 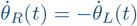.

For a reflector *i* at direction **n**_*i*_, the instantaneous radial velocity of the ear relative to the bat’s head is

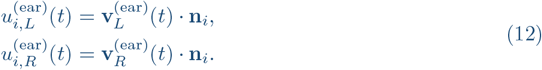

The total effective radial velocity at each ear, combining bat body motion, object motion, and ear motion, becomes

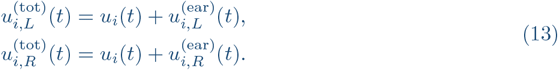

The corresponding echo frequencies at the left and right ears are then

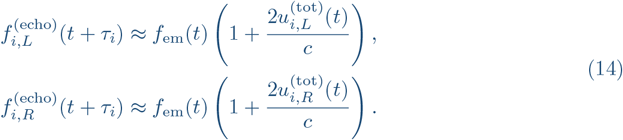

#### 2.2.2 Echo-Only Tagging and Scene Parsing

Crucially, oscillatory ear motion modulates the *echo* but not the outgoing call. The emitted CF tone remains approximately constant at *f*_em_(*t*) during each pulse, whereas the received echo from object *i* at ear *E* ∈ {*L, R*} has a time-varying instantaneous frequency

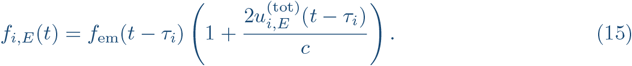

Expressed in terms of instantaneous phase *ϕ*_*i,E*_(*t*), the analytic echo signal at ear *E* can be written as

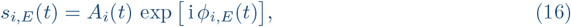

with instantaneous frequency

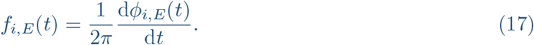

Because 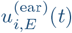 depends on both the direction **n**_*i*_ and the phase of ear motion at echo arrival, each reflector acquires a characteristic *modulation pattern f*_*i,E*_(*t*) and amplitude envelope *A*_*i*_(*t*) at each ear. Two important consequences follow:

1. The outgoing call remains spectrally narrow and only weakly modulated, whereas the echo exhibits sidebands and time-varying frequency due to 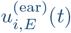. This difference provides an “echo-only” tag that can be exploited to segregate echo from call even when Doppler-shift compensation renders their carrier frequencies nearly identical.
2. Different reflectors *i* and *j* return echoes at different delays *τ*_*i*_, *τ*_*j*_ and at different phases of the ear oscillation cycle, leading to distinct modulation patterns. The binaural disparity

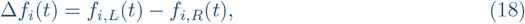

and related quantities (e.g. interaural amplitude differences) thus form an object-specific signature that can help the bat parse the scene and identify potential targets in a cluttered echo stream.

#### 2.2.3 Interaction with Call–Echo Overlap

Let *x*(*t*) denote the emitted CF call of duration *T*_call_ and carrier frequency *f*_em_. At ear *E*, the total received signal is the superposition of the ongoing call (via bone conduction and internal reflections) and the sum of echoes:

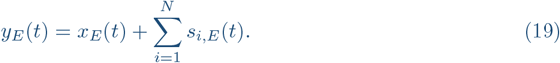

During close-range approach, echo delays *τ*_*i*_ can be shorter than *T*_call_, so that *x*_*E*_(*t*) and *s*_*i,E*_(*t*) overlap in time and frequency. Under global-reference DSC, the carriers of *x*_*E*_ and *s*_*i,E*_ are nearly identical, and classical Doppler separation fails.

Oscillatory ear motion, however, ensures that *s*_*i,E*_(*t*) contains modulation components at and around the ear oscillation frequency *f*_*e*_ (and its harmonics), whereas *x*_*E*_(*t*) does not. By analysing *y*_*E*_(*t*) in a reference frame locked to the known ear motion—for example, by demodulating at *f*_*e*_ or correlating with sin(*ω*_*e*_*t*) and cos(*ω*_*e*_*t*)—the bat can, in principle, recover echo-specific features even under call–echo overlap. In this way, ear oscillations provide a mechanical tagging mechanism that helps separate echoes from the emitted signal.

#### 2.2.4 Range Dependence and the Need for Higher Ear Frequencies

As the bat approaches a potential target at distance *d*(*t*), echo delay *τ* (*t*) ≈ 2*d*(*t*)*/c* and call duration *T*_call_(*t*) both decrease. In CF bats, close-range approach is characterised by shorter calls and high pulse-repetition rates [3, 4]. The effective time window *T*_echo_ during which the echo is available for analysis thus shrinks with decreasing *d*.

Let *T*_echo_ denote the duration over which the echo from a given object is above threshold. To obtain at least *K* distinct cycles of ear-induced binaural contrast within this window, the ear oscillation frequency *f*_*e*_ must satisfy

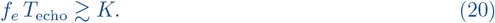

As *d* decreases, *T*_echo_ shrinks, so maintaining a fixed *K* requires

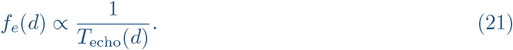

This provides a natural explanation for the empirical observation that ear oscillation rates increase as bats approach a target [3, 4]. Faster ear movements compress multiple cycles of binaural contrast and echo-only modulation into a shorter echo window, stabilising the rate of informative samples under high temporal pressure.

Within this framework, global-reference DSC stabilises echoes from the stationary environment but leaves insect-induced Doppler shifts and call–echo overlap unresolved. Oscillating ears, moving in antiphase, supplement this system by imposing echo-only modulations whose structure depends on object direction, timing, and relative motion. These transformations provide object-specific signatures that facilitate scene parsing during foraging and maintain effective information collection as call duration and echo delay decrease during close-range approach.

### 2.3 Computational Modelling of Distributed Doppler and Phase Transformation

#### 2.3.1 CF Call and Simulation Parameters

A 100-ms constant-frequency call at 80 kHz was synthesised at a sampling rate of 192 kHz to approximate narrowband pulses typical of CF bats. The bat was modelled as flying forward at a constant velocity of 2 m/s in still air (speed of sound: 343 m/s). The simulated echo was computed for binaural receivers (left and right ears), with time-varying motion profiles imposed on each ear in antiphase. (Fig. 2 top panel)

**Figure 2:**
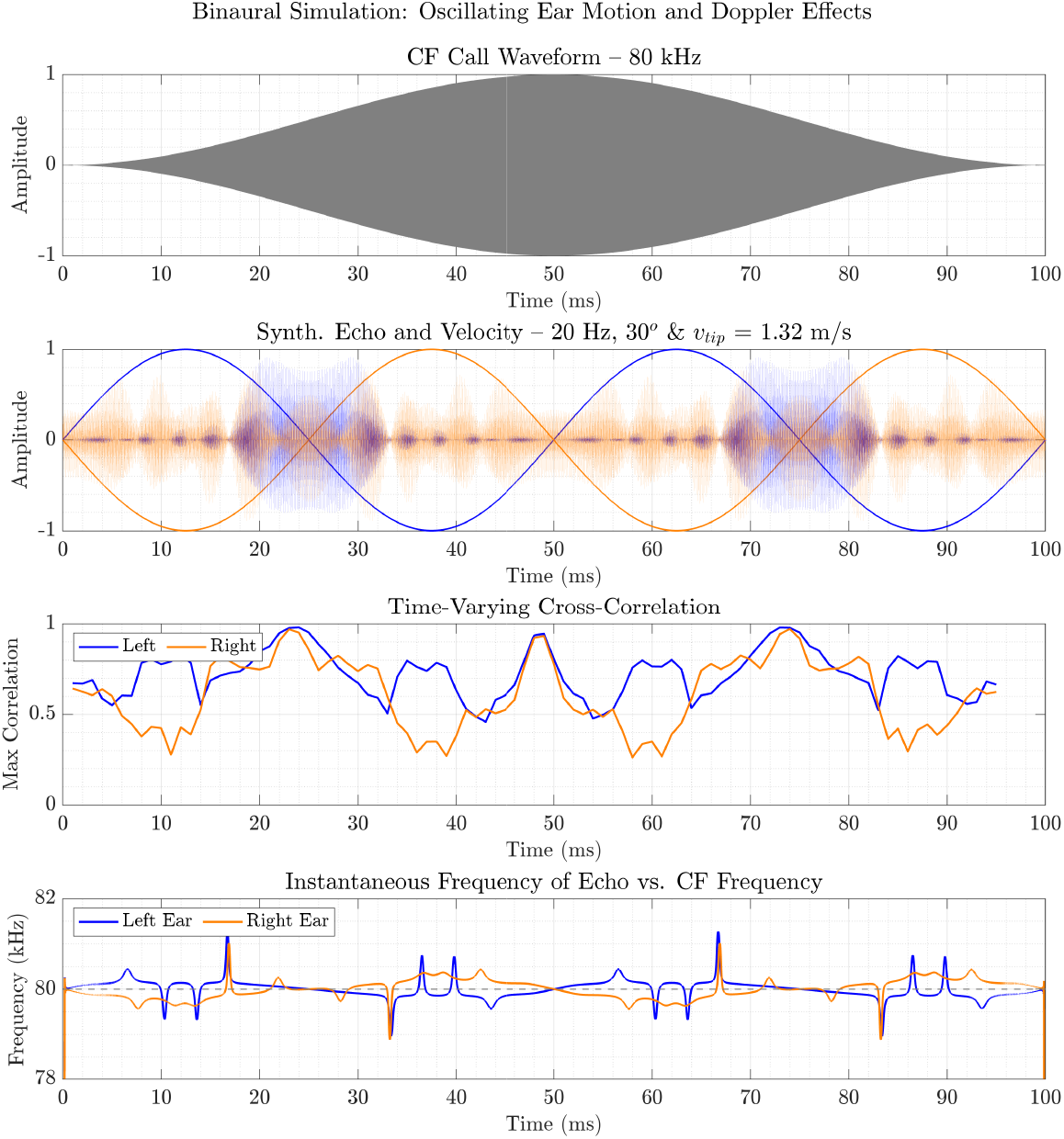
Binaural simulation of echo transformation by oscillating ear motion. This figure illustrates the simulated effect of ear oscillations on binaural echo reception for a constant-frequency (CF) call. The top panel shows the 80 kHz synthetic CF call waveform lasting 100 ms. The second panel displays the simulated echoes received at each ear as translucent coloured patches (left: blue, right: orange), overlaid with the corresponding normalised tip velocity profiles (solid lines). Due to opposing sinusoidal ear motions (20 Hz, ± 30°), the echoes are dynamically warped in both amplitude and phase. The third panel shows the time-varying cross-correlation between the left and right ear signals, revealing dynamic changes in binaural timing alignment induced by phase distortion and frequency shifts. The bottom panel plots the instantaneous frequency in each ear, demonstrating the divergent Doppler-induced modulation imposed by the oscillating ear motion. Together, these panels reveal how even a CF echo is transformed into a spectrotemporally rich, binaurally asymmetric signal through biologically plausible ear kinematics.

#### 2.3.2 Ear Kinematics and Doppler-Shifted Echo Modelling

Each ear was represented as a vertical structure of height 2 cm undergoing angular oscillation with maximum deflections of 15°, 30°, or 45°, and frequencies ranging from 5 to 50 Hz. The peak linear velocity at the ear tip was computed as:

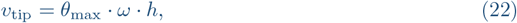

where *ω* = 2*πf*_ear_ is the angular velocity, *θ*_max_ is the angular displacement (in radians), and *h* is the ear height.

The ear was discretised into 100 vertical segments from base to tip. For each segment, the instantaneous linear velocity followed a sinusoidal pattern scaled by height. The Doppler-shifted frequency at each time point was computed as:

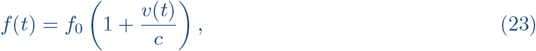

where *f*_0_ is the CF frequency, *v*(*t*) is the segment velocity, and *c* is the speed of sound. The echo was constructed by integrating the instantaneous phase at each time point:

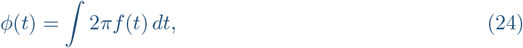

Thus, modelling frequency shifts due to motion as continuous phase warping. Each segment’s contribution was delayed based on its one-way travel time, and the total echo was synthesised by summing across segments. A toggle allowed switching between summation and amplitude-averaging across segments (testing), but all presented results used summation. The methods of the simulation are depicted in Figure 2.

### 2.4 Instantaneous Frequency Estimation

The analytic signal of each echo was computed via the Hilbert transform. Instantaneous frequency was obtained from the temporal derivative of the unwrapped phase:

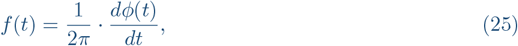

where *ϕ*(*t*) = arg[*z*(*t*)] and 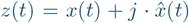. Regions with envelope magnitudes below 1% of the maximum were excluded, and the frequency trace was smoothed using a 50-sample moving average. Only the central 60 ms of each trace (excluding the first and last 20 ms) was retained for analysis to avoid onset and offset artefacts (Fig. 2 bottom panel).

#### 2.4.1 Spectrogram and Visualization

Time–frequency representations were obtained via short-time Fourier transforms using a 2048sample Hamming window and 95% overlap. Spectrogram overlays were constructed with grayscale encoding for the left ear and red channel encoding for the right ear. Other panels included waveforms, ear tip velocity profiles, and plots of Δ*f* (*t*) and BLD with dual y-axes and consistent colour themes. (Fig. 4)

**Figure 3:**
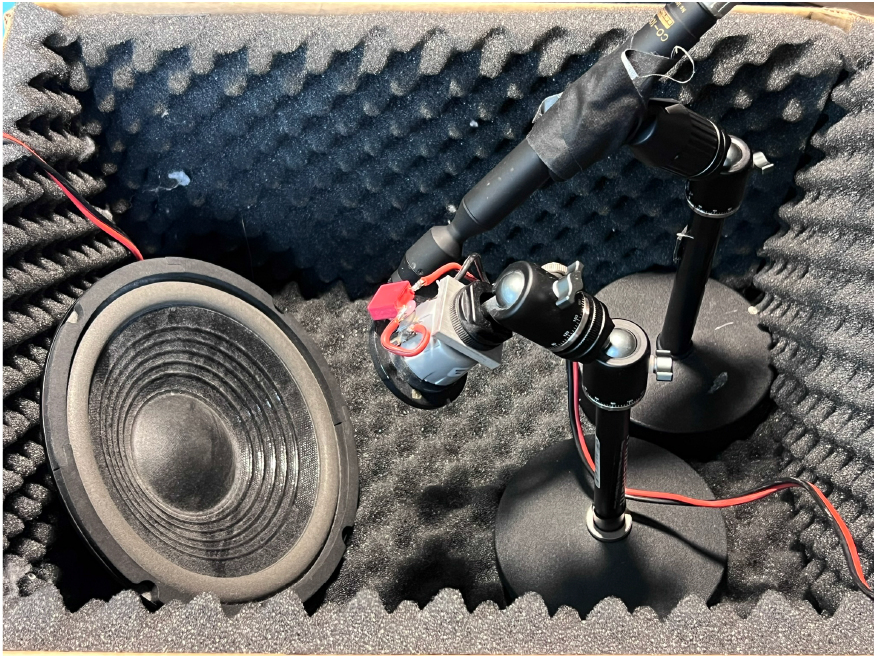
Experimental setup for measuring motion-induced Doppler effects. A custom-built ESP32-based playback system was used to generate oscillations in a 20 cm subwoofer via an I^2^S DAC (PCM5102) and a 50 W Class D amplifier (ZK-TB21). Oscillatory waveforms at 5–30 Hz were created in MATLAB, saved as .wav files, and played back from a microSD card using custom firmware flashed via PlatformIO. An 80 kHz tone was generated and recorded using an RME Babyface Pro interface. A Sanken CO-100K microphone was aligned with the tone playing speaker (Scanspeak, R2004), both directed at the centre of the subwoofer, positioned 20 cm away. Control trials were recorded with the subwoofer inactive.

**Figure 4:**
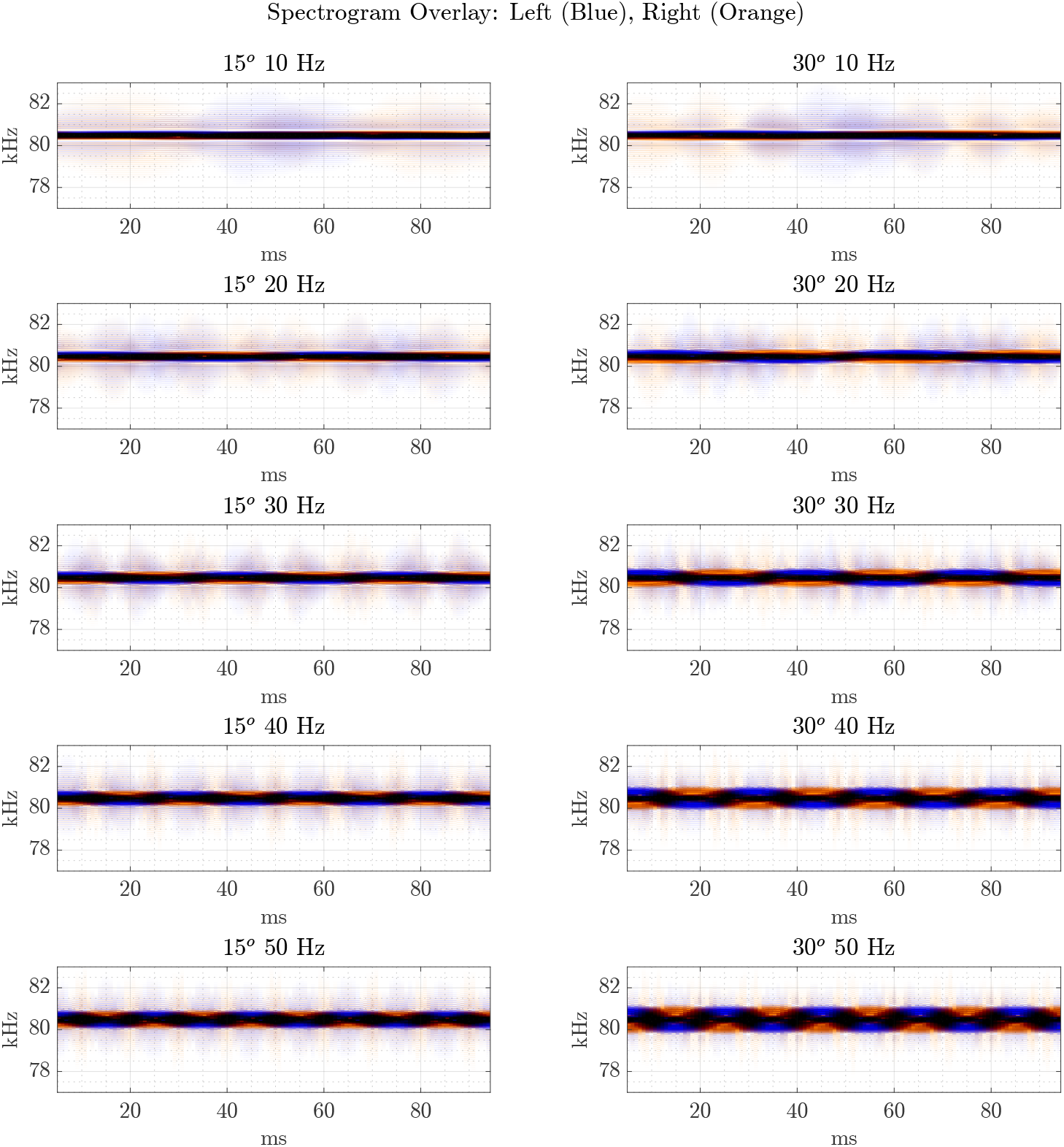
Spectrogram overlays of binaural echoes for varying ear oscillation conditions. Each subplot shows time–frequency representations of echoes received at the left (blue) and right (orange) ears under different angular displacements (15°, 30°) and ear oscillation frequencies (10–50 Hz). As oscillation frequency and angular displacement increase, spectral modulation deepens and broadens, producing pronounced dynamic variation around the 80 kHz CF tone.

#### 2.4.2 Parameter Sweep and Output Summary

Simulations were conducted for 18 combinations of angular frequency (5–50 Hz) and displacement (15–45°). For each run, the resulting echo metrics and figures were saved. The peak tip velocity, Δ*f* bandwidth, standard deviation of Δ*f*, and BLD were collected in a summary table. The full output was saved as CSV and MATLAB ‘.mat’ files. (Fig. 6)

**Figure 5:**
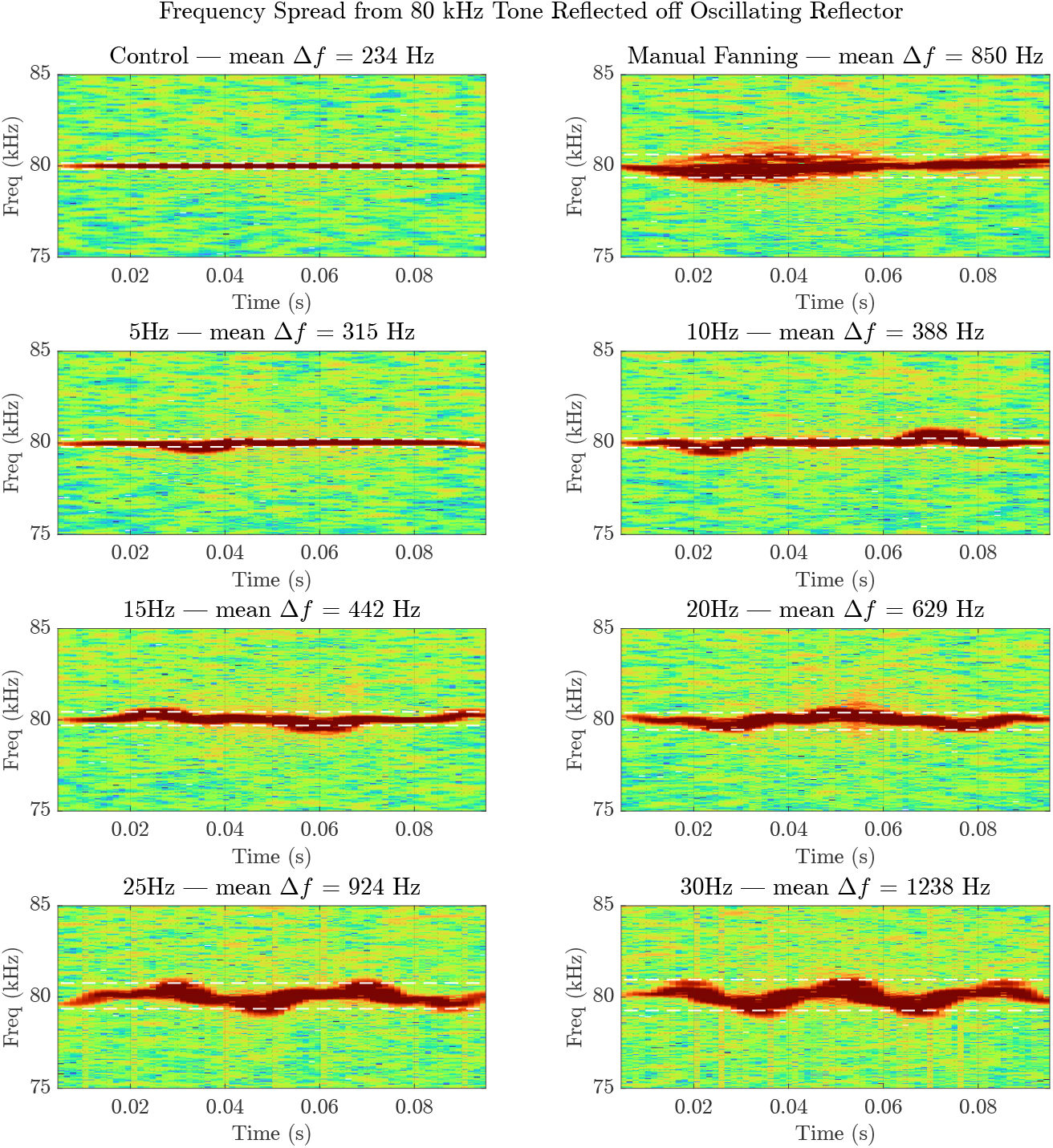
Spectrograms of reflected 80 kHz tone showing frequency spread across conditions. Each panel displays the time–frequency representation of a 100 ms segment extracted from the reflected signal for the corresponding trial condition. The analysis was performed using short-time Fourier transform with a window length of 2048 samples and a 90% overlap. Dashed white lines indicate the upper and lower frequency boundaries of the tone’s spectral energy within the **-20 dB** band relative to its peak power. The numeric label in each title represents the mean Δ*f* (frequency spread) across all segments in that condition. In the **Control** condition, the spectrogram shows a narrowband, nearly stationary tone centred at 80 kHz, with minimal frequency spread (mean Δ*f* = 234 Hz). As reflector motion increases (from 5 Hz to 30 Hz), the signal exhibits progressively broader frequency modulation, with Δ*f* increasing accordingly. This spread reflects the Doppler effect induced by oscillatory motion, which introduces instantaneous frequency shifts. At higher modulation frequencies (e.g., 25 Hz and 30 Hz), the bandwidth widens considerably, indicating more pronounced phase distortion of the reflected wave. The **Manual Fanning** condition also produces a large frequency spread, though with more irregular temporal structure, likely due to non-sinusoidal, variable-speed motion. These results indicate what may be expected from oscillating ears, with increasing oscillation frequency leading to greater spectral broadening of the reflected signal, consistent with motion-induced Doppler dynamics.

**Figure 6:**
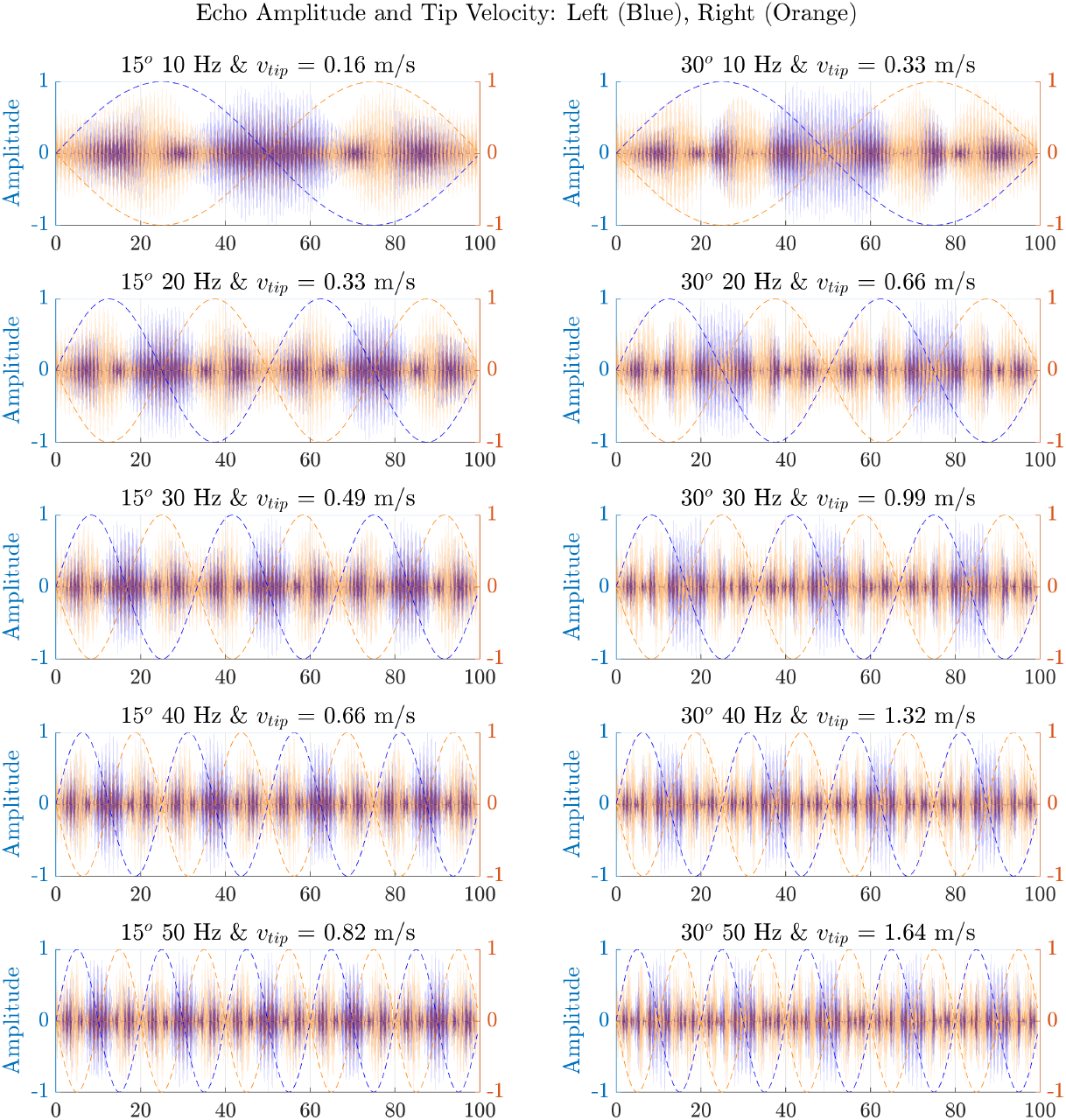
Simulated echo waveforms and corresponding ear tip velocity profiles. Left and right ear echoes are plotted alongside their respective velocity traces for multiple ear oscillation frequencies and angular displacements. Echo amplitudes and timing shift with velocity phase, illustrating dynamic modulation of the received signal imposed by oscillatory motion.

##### Binaural Frequency and Level Differences

Binaural frequency disparity was computed as:

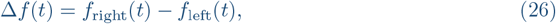

and binaural level difference (BLD) as:

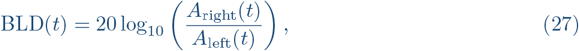

where *A*(*t*) denotes the envelope of the analytic signal. The standard deviation of both Δ*f* and BLD was computed from the valid region. The Δ*f* bandwidth was defined as the maximum of |Δ*f* (*t*)| within the selected segment. (Fig. 5)

#### 2.4.3 Implementation

All simulations and post-processing were conducted using MATLAB R2023a. Custom scripts handled the synthesis of CF calls, Doppler and phase transformations, segment-wise integration, and Hilbert-based feature extraction.

### 2.5 Experimental Approach

#### 2.5.1 Oscillation Playback System

I developed a custom tone modulation system using an ESP32 microcontroller connected to a 20 cm subwoofer. The ESP32 was interfaced with a PCM5102 I^2^S DAC for digital audio output, which was amplified via a 50 W Class D amplifier module (model ZK-TB21, China). The ESP32 firmware was developed and compiled using PlatformIO and flashed to the board over USB. This firmware was programmed to read and play back .wav files stored on a microSD card. Oscillation waveforms were generated in MATLAB as pure sine tones with frequencies ranging from 5 to 30 Hz, sampled at 44.1 kHz, and saved in 16-bit .wav format. These tones were looped in sequence by the ESP32, driving sinusoidal motion in the subwoofer cone (Fig. 3). The experimental setup is not intended to reproduce pinna geometry or directional filtering, but to isolate the acoustic consequences of relative oscillatory motion under controlled conditions.

#### 2.5.2 Recording Configuration

Recordings were made using a Sanken CO-100K microphone aligned parallel to the Scanspeak tweeter (Model R2004, Scan-Speak A/S, Herning, Denmark) at a distance of 20 cm, both aimed at the centre of the subwoofer membrane. The microphone signal was routed through an RME Babyface Pro (RME GmbH, Haimhausen, Germany) audio interface operating at 192 kHz. A MATLAB script was used to generate a series of 100 ms long 80 kHz tones with 1-second intervals. These tones were simultaneously played back and recorded in MATLAB using the RME interface. An internal trigger loopback signal was recorded on the second channel to assist signal segmentation.

To synchronise playback with the subwoofer oscillation, the MATLAB recording script was manually triggered immediately after ESP32 playback began. Each trial consisted of 10 repetitions of the tone, and the entire sequence was recorded for analysis. Control recordings were taken at the end of the session with the subwoofer powered off to assess baseline frequency spread without motion.

#### 2.5.3 Spectral Analysis

Recorded signals were segmented based on a synchronisation trigger embedded in the second channel. Each 100 ms segment was analysed using a short-time Fourier transform with a 2048-sample window, 1800-sample overlap, and a 4096-point FFT. The frequency spread (Δ*f*) of each segment was defined as the bandwidth containing spectral energy within 20 dB of the peak, constrained to the range 75–85 kHz. The Doppler-induced broadening was quantified by comparing Δ*f* across conditions (Fig. 5).

## 3 RESULTS

To evaluate the effects of oscillating receiver motion on the acoustic structure of returning echoes, I conducted both numerical simulations and controlled acoustic experiments. The simulations modelled the reception of CF echoes by antiphase-oscillating ears over a range of kinematic conditions, systematically varying oscillation frequency and angular displacement. In parallel, I performed physical measurements using a stationary speaker and microphone with an oscillating reflector to emulate the relative motion of the bat’s pinna. Together, these two approaches provide converging insights into how dynamic receiver motion transforms echo features, particularly in terms of frequency bandwidth, amplitude envelope, and binaural disparity. The following sections present the results of these simulations and experiments, illustrating their alignment and divergence across different conditions.

### 3.1 Origin and Characteristics of Spectral Spread

Unlike classical Doppler shifts resulting from linear flight, where a fixed relative velocity induces a single frequency offset, oscillatory motion introduces a dynamic velocity profile throughout each echo interval. Because CF bat calls can last 50–100 ms, and the echo arrives while the ear is in motion, the reflected signal experiences time-varying phase shifts. These result in not just a shift, but a spectral broadening—a band of frequencies reflecting the changing instantaneous velocity of the reflector.

This effect is fundamentally different from the classical Doppler shift caused by constant linear velocity. In the case of a bat flying steadily at 3 m/s, the relative velocity between source and receiver remains fixed. The Doppler effect in such a scenario causes a uniform frequency shift given by the well-known relation:

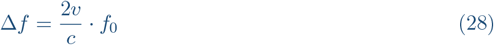

For a tone of 80 kHz and *v* = 3 m/s, this yields a constant Doppler shift of approximately 1400 Hz. Importantly, *the entire frequency spectrum is translated*, and the output remains spectrally narrow–only offset in frequency. There is no spread, modulation, or temporal variation in spectral content.

By contrast, when a bat’s ear oscillates sinusoidally, the velocity of the reflective surface (i.e., the ear) is continuously changing throughout the echo duration. The instantaneous velocity *v*(*t*) varies as a sinusoidal function:

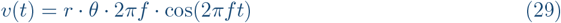

where *r* is the length from the base to the ear tip, *θ* is the angular amplitude in radians, and *f* is the ear oscillation frequency. This variation in *v*(*t*) results in a continuously changing instantaneous Doppler shift, rather than a constant one. The received echo is thus not simply shifted, but frequency modulated.

Mathematically, this is equivalent to applying a time-varying phase term to the reflected signal. The result is not a displaced tone, but a signal whose spectrum is spread across a range of frequencies. This manifests as spectral broadening (bandwidth spread), which depends on the velocity range of the ear tip during the echo interval.

Table 1 summarises the outcomes of 30 binaural Doppler simulations, systematically varying ear oscillation frequency and angular amplitude. The tip velocity, computed from ear kinematics, ranged from 0.08 to 2.47 m/s, corresponding to angular excursions of 15° to 45° and frequencies from 5 to 50 Hz. The resulting binaural frequency bandwidths (Δ*f*) showed a strong positive trend with increasing tip velocity, rising from 107 Hz at 5 Hz, 15° to over 1360 Hz at 50 Hz, 45°. The standard deviation of Δ*f* also scaled with velocity, indicating that higher oscillation amplitudes not only broadened the frequency spread but introduced greater temporal variability. Interestingly, across all conditions, the binaural level difference (BLD) standard deviation remained relatively stable, clustering around 6.2 dB at peak excursions. These results confirm that rapid angular ear motion introduces dynamic frequency shifts and binaural disparities of biologically significant magnitude, consistent with the hypotheses on their functional importance in echolocating CF bats.

**Table 1:** Summary of simulated binaural Doppler conditions across combinations of ear oscillation frequency and angular excursion. Each trial reports the resulting tip velocity and corresponding tip displacement (Excursion), along with the peak binaural frequency disparity (Δ*f* bandwidth), its standard deviation, and the standard deviation of the binaural level difference (BLD). All simulations assume a constant forward flight velocity of 2 m/s and a 20 mm ear length.

| Trial | Ear Freq(Hz) | Ang.(°) | Tip $v$ (m/s) | Excur.(mm) | $\Delta f$ BW(Hz) | $\Delta f$ $\sigma$(Hz) | BLD $\sigma$(dB) |
| --- | --- | --- | --- | --- | --- | --- | --- |
| 2 | 5 | 15 | 0.08 | 5.2 | 23.1 | 8.8 | 3.6 |
| 3 | 5 | 30 | 0.16 | 10.5 | 67.9 | 30.9 | 5.4 |
| 4 | 5 | 45 | 0.25 | 15.7 | 113.0 | 43.6 | 6.2 |
| 5 | 10 | 15 | 0.16 | 5.2 | 107.4 | 65.4 | 3.6 |
| 6 | 10 | 30 | 0.33 | 10.5 | 201.8 | 95.1 | 5.5 |
| 7 | 10 | 45 | 0.49 | 15.7 | 213.2 | 109.9 | 6.2 |
| 8 | 15 | 15 | 0.25 | 5.2 | 160.8 | 83.2 | 3.6 |
| 9 | 15 | 30 | 0.49 | 10.5 | 318.8 | 129.8 | 5.4 |
| 10 | 15 | 45 | 0.74 | 15.7 | 420.7 | 187.6 | 6.2 |
| 11 | 20 | 15 | 0.33 | 5.2 | 214.1 | 107.8 | 3.6 |
| 12 | 20 | 30 | 0.66 | 10.5 | 423.7 | 179.2 | 5.5 |
| 13 | 20 | 45 | 0.99 | 15.7 | 558.8 | 236.7 | 6.2 |
| 14 | 25 | 15 | 0.41 | 5.2 | 267.4 | 139.0 | 3.6 |
| 15 | 25 | 30 | 0.82 | 10.5 | 528.4 | 232.7 | 5.4 |
| 16 | 25 | 45 | 1.23 | 15.7 | 695.9 | 309.4 | 6.2 |
| 17 | 30 | 15 | 0.49 | 5.2 | 320.3 | 182.0 | 3.6 |
| 18 | 30 | 30 | 0.99 | 10.5 | 631.7 | 301.9 | 5.5 |
| 19 | 30 | 45 | 1.48 | 15.7 | 831.2 | 400.8 | 6.2 |
| 20 | 35 | 15 | 0.58 | 5.2 | 373.1 | 206.6 | 3.6 |
| 21 | 35 | 30 | 1.15 | 10.5 | 734.5 | 340.2 | 5.4 |
| 22 | 35 | 45 | 1.73 | 15.7 | 965.8 | 446.8 | 6.2 |
| 23 | 40 | 15 | 0.66 | 5.2 | 425.6 | 224.1 | 3.6 |
| 24 | 40 | 30 | 1.32 | 10.5 | 836.6 | 381.4 | 5.5 |
| 25 | 40 | 45 | 1.97 | 15.7 | 1099.0 | 521.2 | 6.2 |
| 26 | 45 | 15 | 0.74 | 5.2 | 477.9 | 251.6 | 3.6 |
| 27 | 45 | 30 | 1.48 | 10.5 | 937.5 | 417.1 | 5.4 |
| 28 | 45 | 45 | 2.22 | 15.7 | 1231.7 | 547.8 | 6.2 |
| 29 | 50 | 15 | 0.82 | 5.2 | 530.1 | 290.8 | 3.6 |
| 30 | 50 | 30 | 1.64 | 10.5 | 1038.2 | 480.0 | 5.5 |
| 31 | 50 | 45 | 2.47 | 15.7 | 1364.2 | 629.4 | 6.2 |

In my experiments, reflector oscillation at 30 Hz with *r* = 2 cm and *θ* = 45° produced instantaneous velocities up to 2.96 m/s (Calculated based on speaker parameters). This yielded a dynamic frequency excursion of approximately 1.2 kHz around the 80 kHz carrier (Figure 5). Unlike the constant offset from forward motion, this spread is transient, phase-locked to the modulator, and results in temporally rich echo transformations. This modulation is further reflected in the waveform domain as phase warping, causing irregular zero-crossings and beat-like structures–features entirely absent in linear Doppler conditions (Fig. 7).

**Figure 7:**
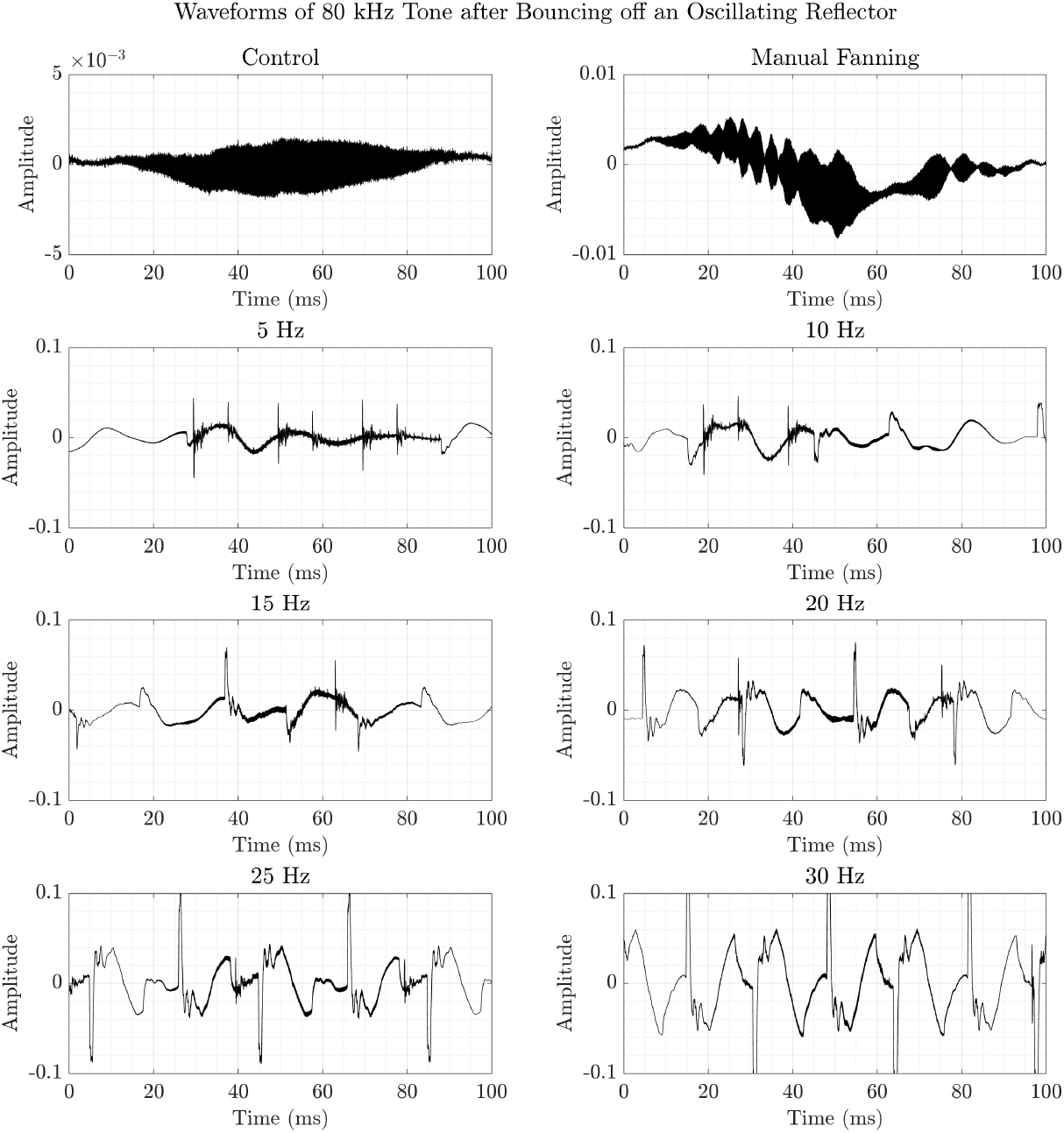
Time-domain waveforms of 80 kHz tone after reflection from an oscillating surface. Each panel shows a 100 ms segment extracted from the recorded signal for the corresponding condition. The **Control** condition (top left) reveals a clean and stable sinusoidal waveform, characteristic of a continuous tone with no motion-induced modulation. In contrast, all other conditions introduce varying degrees of non-stationarity due to surface movement. At low modulation rates (5–10 Hz), the reflected waveform exhibits periodic amplitude fluctuations and mild distortions. As the oscillation frequency increases (15–30 Hz), the waveform becomes increasingly distorted, with prominent changes in both amplitude and zero-crossing structure. This reflects **phase warping** caused by the Doppler shift imparted by the moving reflector. Notably, in the higher frequency conditions (especially 25 Hz and 30 Hz), the waveform appears stretched and compressed in a quasi-periodic manner, producing variable intervals between zero-crossings–unlike the uniform spacing seen in the Control. Manual fanning yields a complex, irregular pattern resembling fluctuating velocity components, which also alters zero-crossing symmetry. These features demonstrate how relative motion between emitter and reflector induces instantaneous frequency modulation and waveform distortion even in a nominally constant-frequency tone.

These findings demonstrate that ear oscillations act as a signal transformation mechanism, not just a positional adjustment. They dynamically encode time-varying velocity and phase, and cannot be simplified to a static DS model. Therefore, the acoustic consequence of oscillating ears in CF bats must be interpreted as *Doppler-based modulation*, not just shift, and forms a biologically distinct strategy for echo encoding.

I systematically simulated the effects of oscillatory ear motion on echo reception across a range of ear oscillation frequencies (5–50 Hz) and angular displacements (15°, 30°, 45°). The simulations produced a set of echo waveforms for the left and right ears, which were analysed in terms of spectral characteristics (Fig. 4), waveform and velocity dynamics (Fig. 6), and binaural differences in frequency and amplitude (Fig. 8).

**Figure 8:**
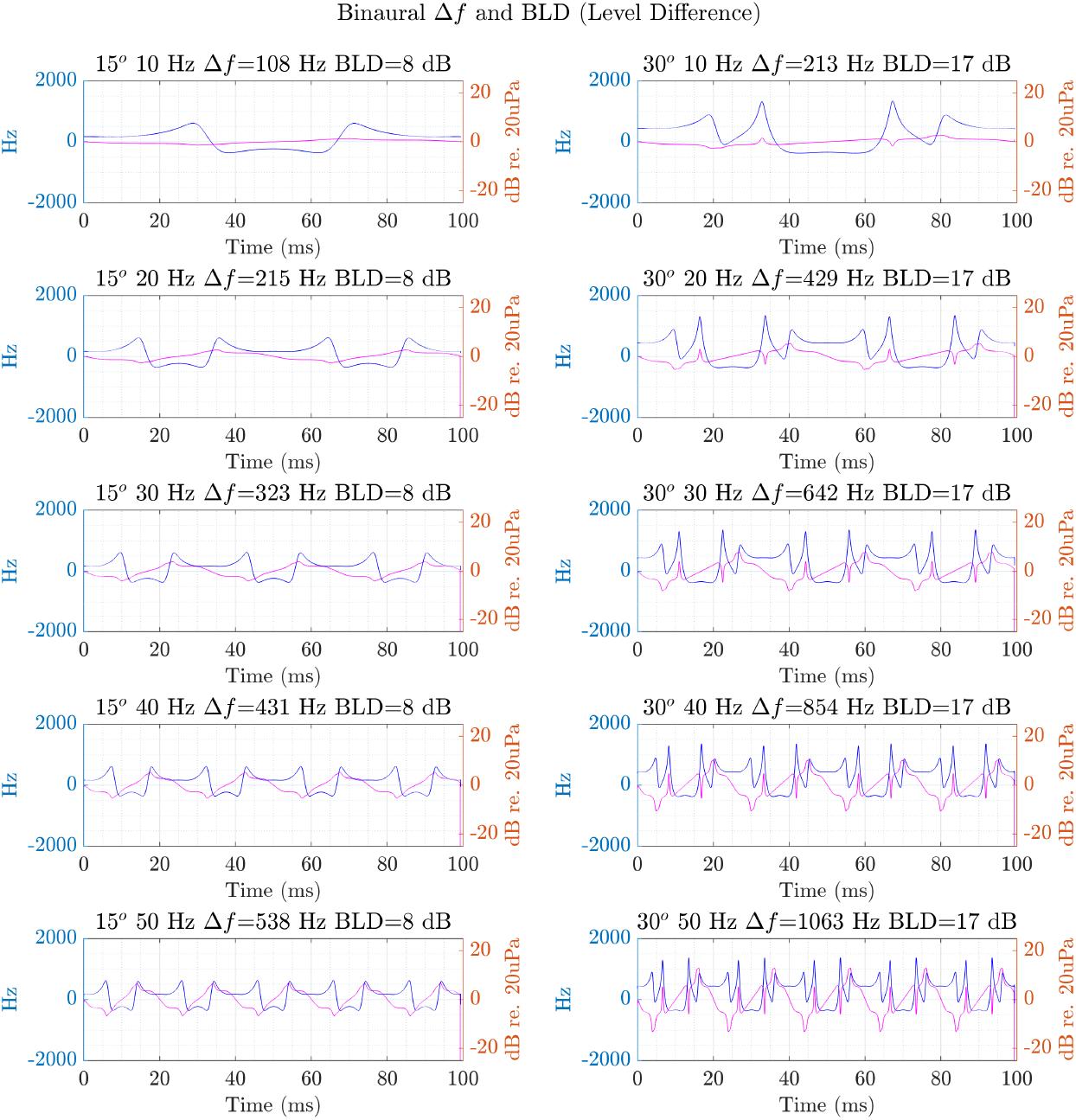
Instantaneous binaural frequency difference (Δ*f*) (*magenta*) and binaural level difference (BLD) (*blue*) traces for each simulation condition. Plots show how dynamic ear motion produces non-linear binaural disparities. Higher oscillation frequencies and angular displacements lead to stronger modulations in both frequency and amplitude, highlighting the potential role of ear kinematics in encoding spatial acoustic cues.

### 3.2 Spectral Modulation by Ear Motion

Figure 4 shows the spectrograms of the left (blue) and right (orange) ear echoes across all simulated conditions. At low oscillation frequencies (e.g., 5–10 Hz) and small angular displacements, the echo remained close to the transmitted constant-frequency (CF) tone at 80 kHz. However, with increasing ear frequency and angular range, I observed progressive spectral broadening and modulation. For instance, at 50 Hz and 45°, the spectral content extended over several hundred Hertz, indicating strong Doppler-induced shifts caused by rapid and large ear movements. Importantly, the left and right spectrograms diverged significantly due to their antiphase motion, with distinct spectral energy contours suggesting differential encoding of spatial information.

Similarly, the spectrograms from the recordings revealed that the frequency spread increased proportionally with oscillation frequency (Fig. 5). Control trials showed narrowband reflections with Δ*f* around 234 Hz. By contrast, 30 Hz oscillations produced spectral spreads approaching 1.3 kHz. These effects are temporally dynamic and phase-locked to the oscillator’s motion. Waveforms also exhibited noticeable phase warping: zero-crossings became irregular, with beat-like modulations arising from interference patterns. Such distortions were absent in the Control and are not predicted by linear Doppler alone.

### 3.3 Temporal Alignment with Ear Velocity

The temporal structure of the echoes was also modulated by ear movement. As shown in Figure 6, the amplitude envelopes of the echoes vary in close synchrony with the sinusoidal velocity profiles of the ear tips. Echo peaks tended to align with peak velocities, while troughs coincided with direction reversals. This dynamic interaction between incoming echoes and moving receiver geometry led to periodic constructive and destructive interference in the waveform. The left and right ears displayed mirror-symmetric patterns, consistent with their opposite motion phases. These effects intensified with higher frequencies and greater angular sweeps. Figure 7 presents the waveforms of the recordings from the oscillating target. The expected phase warping and the resulting zero crossing distortion become increasingly evident with higher oscillation frequencies.

### 3.4 Binaural Differences in Frequency and Level

Figure 8 illustrates the instantaneous binaural frequency difference (Δ*f*) and binaural level difference (BLD) as functions of time. At low oscillation frequencies, both Δ*f* and BLD remained relatively stable and small in magnitude. As ear movement velocity increased, however, Δ*f* fluctuated more rapidly and reached larger extrema. Similarly, BLD variation widened, reflecting the amplitude asymmetry introduced by differing Doppler and phase interactions at each ear. These features represent dynamic binaural cues potentially exploitable by the bat’s auditory system for spatial localisation or motion tracking.

Overall, the results demonstrate that oscillatory pinna motion can impose rich, condition-dependent modulations on received echoes in both spectral and temporal domains. These transformations scale systematically with the biomechanical parameters of ear motion, offering a potential mechanism for sensory enhancement beyond directional gain.

These findings confirm that ear oscillations generate rich transformations to returning echoes– transformations that scale nonlinearly with velocity and encode motion phase. This suggests a role for ear dynamics in enriching spatial and temporal encoding in CF bat biosonar.

## 4 DISCUSSION

This study combined a theoretical framework, computational modelling, and an empirical validation experiment to examine how oscillatory ear movements in constant-frequency (CF) bats transform the structure of returning echoes. By grounding the analysis in the receiver-side Doppler formulation (Eqs. 3, 14) and explicitly modelling the distributed velocity field along an oscillating pinna (Eqs. 14), I show that ear motion produces echo-only spectrotemporal signatures that cannot arise from Doppler-shift compensation (DSC) or static directional filtering. Across theory, simulation, and experiment, a consistent functional interpretation emerges: oscillatory ears act as dynamic acoustic transformers that enhance echo information precisely when conventional Doppler cues become unreliable.

### 4.1 Theoretical Implications: Oscillatory Reception Generates Echo-Only Transformations

The theoretical framework (Section 2.1 & Fig. 2) highlights three key principles.

First, the instantaneous frequency perceived by a moving receiver depends on the receiver’s radial velocity relative to the source (Eq. 3). Because a bat’s ear is an extended, moving structure, each point along the pinna experiences a distinct instantaneous velocity (Eqs. 14), generating a spatially distributed Doppler field across the receiving surface.

Second, each ear segment contributes a phase-accumulated component to the echo, with its own trajectory *ϕ*_*i*_(*t*) determined by its local motion. The composite echo is thus a superposition of many slightly shifted sinusoids, establishing oscillatory reception as a *phase-warping* mechanism rather than a simple frequency shift.

Third, the ears oscillate in antiphase (Fig. 2), creating dynamic binaural differences in instantaneous frequency and amplitude. These disparities are not simple mirror images because nonlinear kinematics and phase accumulation interact. The framework therefore predicts monaural modulation, spectral broadening, and phase-dependent sampling of different reflectors — all emerging directly from receiver motion.

### 4.2 Simulation Results Confirm Dynamic Spectral and Binaural Structure

The simulations (Figs. 4, 6, 8) instantiate these principles with biologically realistic parameters.

#### Instantaneous frequency fluctuations

The predicted 0.5–1 kHz shifts within a single oscillation cycle arise directly from segment-specific velocities and are visible in the instantaneous frequency traces (Fig. 2, bottom panel). These fluctuations fall well within the perceptual resolution of CF bats and provide temporal landmarks for distinguishing echo structure during call–echo overlap.

#### Spectral broadening

Combining phase-warped components yields spectral broadening (Fig. 4) that scales systematically with oscillation velocity. This broadening introduces sidebands around the received CF frequency, increasing the probability that some echo energy falls within the auditory fovea even when the carrier overlaps the outgoing call.

#### Dynamic binaural contrast

Antiphase ear motion generates alternating cycles of strong and weak binaural contrast, manifesting as time-varying binaural frequency differences and level differences (Fig. 8). Because echoes from different objects arrive at different phases of the oscillation, each reflector acquires a characteristic modulation signature.

#### Dependence on oscillation frequency

Higher oscillation rates amplify these transformations and compress more binaural cycles into shorter echo windows (Figs. 4, 6), consistent with behavioural observations that CF bats increase ear-oscillation frequency at close range.

### 4.3 Experimental Validation of Motion-induced Spectral Transformations

The oscillating-reflector experiment (Figs. 3, 5, 7) provides empirical evidence that motion of the receiver produces nonlinear spectral spreading. The observed increase in spectral bandwidth — from ~234 Hz in the control condition to *>* 1.2 kHz at 30 Hz oscillation — matches the magnitudes and scaling predicted by the simulations (Fig. 5). Although the subwoofer does not mimic the geometry of a bat’s ear, its controlled sinusoidal motion confirms the acoustic principle: oscillatory reception imposes frequency and phase modulations that are both large and structured enough to serve as sensory cues.

### 4.4 Behavioural and Physiological Integration

These findings integrate with and extend the known sensory ecology of CF bats. Doppler-shift compensation stabilises echoes from stationary backgrounds but fails when relative velocity drops near zero [1]. Long CF pulses commonly overlap their own returning echoes [7, 8], leaving little intrinsic temporal or spectral structure to support segregation.

Transmitter–receiver matching in *Rhinolophus* spp. [9, 10] ensures optimal flutter sensitivity but cannot generate additional structure when overlap becomes severe. The echo-only modulations produced by oscillatory reception therefore fill a perceptual gap that neither the transmitter nor the cochlear fovea can resolve.

Biomimetic studies by Yin and Müller [5, 11] demonstrate that moving pinnae generate Doppler-derived directional signatures. However, those models do not incorporate call–echo overlap, antiphase oscillation, or the collapse of object-specific Doppler cues during approach. The present findings therefore complement these studies by identifying a functional role for oscillatory reception specifically under high-duty-cycle conditions unique to CF bats.

### 4.5 Dynamic Binaural Processing and Naturalistic Scene Analysis

Bats exploit extremely small interaural level and phase differences, as well as time-varying spectral cues, to determine object position in three dimensions. Neural recordings in free-flying bats show that spatial representations update dynamically across call–echo cycles [12], while studies of naturalistic stimuli demonstrate sensitivity to echo history, clutter, and multiple simultaneous reflectors [13, 14]. Such processing demands cues beyond delay and mean frequency alone.

Active control of reception is well documented. Head and pinna movements enhance acoustic cues for 3-D localisation [15], and CF bats belong to a guild specialised for detecting micro-Doppler from fluttering targets [16]. Binaural information also constrains DSC stability [17, 18], underscoring how strongly CF bats rely on delicate interaural differences.

Bats can also use object-specific spectral patterns for classification [19] and integrate spatial–temporal cues to discriminate object width and layout [20]. In this context, the modelled oscillation-induced disparities provide a plausible physical basis for “glimpses” of individual reflectors: because each object is sampled at a different oscillation phase, its echo acquires a distinct modulation pattern that can be separated from others and tracked over time.

Robotics-based analyses further support this interpretation. Walker et al. (1998) [21] demonstrated that a mobile robot equipped with two microphones could recover three-dimensional source location from a single constant-frequency tone by exploiting motion-induced binaural disparities. In the framework presented here, oscillatory ear motion plays an analogous role: instead of whole-body movement, the pinnae themselves generate structured, time-varying binaural cues, compressing spatial inference into rapid, biologically feasible ear oscillations.

### 4.6 Temporal and Phase Sensitivity in CF Bats

Further support for oscillatory reception comes from the high temporal and phase sensitivity documented in bat auditory systems. Behavioural studies show that bats can detect sub-microsecond variations in echo timing [22, 23]. Neurophysiological work demonstrates sensitivity to interaural time differences in the envelopes of high-frequency sounds [24, 25], providing a mechanism for exploiting phase-derived cues even without carrier phase locking. In CF bats specifically, neurons in the inferior colliculus and auditory cortex respond strongly to dynamic binaural cues during simulated acoustic motion [26], and the midbrain exhibits an expanded representation of the Doppler-compensated frequency [27]. These specialisations position the auditory system to exploit the instantaneous frequency and binaural envelope structure generated by oscillatory reception.

### 4.7 Dynamic Head-Related Transfer Functions and Oscillatory Reception

The transformations described here can be situated naturally within the head-related transfer function (HRTF) framework by treating the pinnae not as static spectral filters but as *timevarying* ones. In classical terms, the HRTF summarises how head and external-ear geometry impose direction-dependent gains and phase delays at the tympanum; in bats, these directionality functions can be highly frequency specific and behaviourally consequential. In CF/FM echolocators, hearing directionality around the relevant ultrasonic bands has been quantified and shown to depend strongly on external-ear geometry and frequency, providing a mechanistic basis for binaural cue formation even at high carrier frequencies [28]. In CF bats, however, the long, narrowband nature of the carrier means that a *static* HRTF offers only a limited snapshot of direction-dependent filtering over the brief time window in which call–echo overlap and clutter complicate segregation. Oscillatory reception extends this concept: pinna motion converts a static HRTF into a *dynamic HRTF* whose gain and phase are modulated on the timescale of tens of milliseconds, injecting structured, ear-phase-locked fluctuations into the echo while leaving the emitted call largely unchanged.

Empirical and biomimetic work already supports the premise that bat pinna motion and deformation can rapidly reshape the effective acoustic transfer to the ear. For example, modelling and measurements show that moving pinnae can synthetically alter the receiver’s directional selectivity across harmonics, implying that pinna morphology and motion are jointly tuned to extract spatial information from narrowband echoes [29]. Beyond rigid motion, deformation itself can operate as a fast control variable: ear deformations have been proposed to provide a physical mechanism for rapid adaptation of ultrasonic beam patterns, demonstrating that small morphological changes can translate into sizeable changes in acoustic filtering on behavioural timescales [30]. In the context of the present framework, these studies motivate a concrete interpretation: antiphase oscillations implement an internal, periodic sweep of the receiver-side transfer function, so that different reflectors are sampled under systematically different instantaneous HRTFs (different instantaneous gain/phase states), thereby converting a spectrally sparse CF echo stream into a temporally structured sequence of HRTF-tagged “glimpses” that can support scene parsing under overlap and clutter.

### 4.8 Receiver-side Modulations and their Relation to Echo Glints

In echolocating bats, particularly constant-frequency (CF) species, echo structure is often shaped not only by the receiver but also by the target itself. A central concept in this context is that of *echo glints*: discrete highlights within an echo arising from multiple scattering centres on a target, such as an insect’s body and wings. In CF bats, fluttering insect wings introduce periodic amplitude and phase modulations (“flutter echoes”) [31–33] that are highly salient within the auditory fovea and form the basis of classic flutter-detection strategies [16, 34, 35]. Computational and behavioural analyses have further shown that horseshoe bats can localise prey using dominant glints within such multi-component echoes, even when individual reflections overlap in time [36].

The existence of glints makes it clear that structured echo modulations are not, in themselves, novel. However, glints are fundamentally *target-generated* phenomena: their presence, timing, and strength depend on prey morphology, orientation, and wing kinematics. Crucially, glint structure is therefore intermittent and unreliable under many natural conditions, including clutter, unfavourable aspect angles, or when call–echo overlap masks weak reflections. The present framework addresses a complementary and previously underexplored mechanism: *receiver-generated* echo structure arising from oscillatory ear motion. Unlike glints, oscillation-induced modulations are imposed deterministically by the bat’s own ear kinematics and are present even for spectrally simple reflectors. Because these modulations affect the echo but not the emitted call, they provide an “echo-only” signature that persists under Doppler-shift compensation and during severe call–echo overlap.

A key new insight emerging from this perspective is that oscillatory ear motion can interact constructively with glint-based cues rather than merely duplicating them. Multi-glint echoes arrive as a sequence of delayed components; when the ears oscillate in antiphase, each glint is sampled at a different phase of the ear-motion cycle and thus acquires a distinct receiver-imposed modulation state. This effectively *phase-tags* successive glints with ear-kinematic information, transforming a glint train into a temporally structured sequence of binaural and spectrotemporal *snapshots*. Such tagging could stabilise target localisation and individuation when target-generated flutter cues are weak or ambiguous, and provides a principled mechanism by which CF bats may parse complex echo scenes under overlap and clutter. In this sense, oscillatory reception does not replace glint processing, but extends it by embedding target reflections within a dynamic, internally controlled reference frame defined by ear motion.

## 5 CONCLUSIONS

This study identifies oscillatory ear motion in CF bats as a receiver-side mechanism capable of imposing echo-only spectrotemporal and binaural transformations that are absent from the emitted call. By grounding the analysis in receiver Doppler theory and explicitly modelling the distributed velocity field of an oscillating pinna, I show that ear motion can enrich otherwise narrowband CF echoes with dynamic phase, frequency, and amplitude structure precisely under conditions where classical Doppler cues and call–echo separation become unreliable. The convergence of theoretical analysis, computational modelling, and empirical validation supports the view that oscillatory ears function as active acoustic transformers rather than passive directional filters.

The framework makes several concrete and experimentally testable predictions. A primary prediction is that CF bats should be behaviourally sensitive to modulation components aligned with their own ear oscillation frequency and phase, because these modulations are phase-locked to ear kinematics but absent from the outgoing call. Behavioural experiments using virtual echo paradigms could directly test this by imposing controlled frequency or phase modulations on echoes that either match or violate natural ear-motion statistics. At the neural level, the model predicts modulation of instantaneous frequency, envelope phase, and binaural disparity at rates tied to ear motion rather than to call repetition alone. Simultaneous measurements of neural activity and ear kinematics could therefore test whether echo-evoked responses are phase-locked to ear oscillations, particularly within the acoustic fovea. Finally, closed-loop virtual echo experiments that selectively remove oscillation-dependent modulations while preserving overall echo energy offer a direct causal test of whether oscillatory reception contributes information beyond static directional filtering or Doppler-shift compensation.

Together, these predictions position oscillatory ear motion as a biologically plausible solution to the sensory challenges imposed by long-duration CF calls, call–echo overlap, and cluttered acoustic scenes, and provide a path for further validating oscillatory reception as an integral component of CF bat biosonar.

## REVISION SUMMARY

This revised version of *Oscillating Ears Dynamically Transform Echoes in Constant-Frequency Bats* represents a substantial restructuring and strengthening of the original manuscript in response to detailed peer review. The revision introduces a formal theoretical framework that explicitly derives receiver-side Doppler and phase transformations generated by oscillatory ear motion, including distributed velocity fields along the pinna and antiphase binaural effects. These analytical deductions were absent from the initial version and now provide a rigorous physical basis for the proposed mechanism.

The Introduction and Discussion have been rewritten to clarify the specific biological problem addressed — echo processing under call–echo overlap during constant-frequency echolocation — and to situate the work within the broader literature on Doppler-shift compensation, flutter detection, glints, ear motion, and head-related transfer functions. Previously underrepresented and missing prior work has been integrated, and the novelty of the study is now framed explicitly in terms of echo-only, phase-dependent transformations that persist when classical Doppler cues collapse.

The simulation framework has been clarified and the experimental component has been reframed as a controlled physical validation of the underlying acoustic principle rather than a morphological replica of bat pinnae, with methods, assumptions, and limitations stated explicitly.

Overall, the revised manuscript now presents a coherent progression from theory to simulation to experimental validation, offers a clear functional interpretation of oscillatory reception in CF bats, and provides a transparent foundation for future morphology-specific, behavioural, and neurophysiological investigations.

## ACKNOWLEDGEMENTS

I extend my gratitude to colleagues at the Technical University of Munich (TUM) for their valuable discussions and constructive feedback throughout the course of this work. I also thank the three anonymous reviewers for their diligent reading of the original manuscript and for engaging critically with the ideas presented. Their thoughtful comments and careful scrutiny substantially improved the clarity, scope and scientific depth of the revised work, and helped shape the study into its current form.

## COMPETING INTERESTS

The author declares no competing interests.

## FUNDING

This work was conducted without external funding.

## DATA AND CODE AVAILIBILITY

The simulation code, experimental data, and analysis scripts have been publically released on Zenodo DOI: https://doi.org/10.5281/zenodo.15847336.

The corresponding GitHub repository can be accessed via: https://github.com/raviumadi/oscillating_ears.

## Notes

### Competing Interest Statement

The authors have declared no competing interest.

### Summary of Updates

This version introduces a formal receiver-side theoretical framework, clarifies the functional role of oscillatory ear motion under call--echo overlap, integrates relevant prior literature, and strengthens the link between theory, simulation, and experiment. See the end of the manuscript for a more detailed revision summary

